# Sexual maturation and embryonic development in octopus: use of energy and antioxidant defence mechanisms using *Octopus mimus* as a model

**DOI:** 10.1101/310664

**Authors:** Alberto Olivares, Gabriela Rodríguez-Fuentes, Maite Mascaró, Ariadna Sánchez, Karen Ortega, Claudia Caamal-Monsreal, Nelly Tremblay, Carlos Rosas

## Abstract

Sexual maturation and reproduction influence the status of a number of physiological processes and consequently the ecology and behaviour of cephalopods. Using *Octopus mimus* as model species, the present study examined the changes in biochemical composition that take place during gonadal maturation of octopus females and its consequences in embryo and hatchlings characteristics, including energetic metabolites, digestive enzymes and antioxidant defence mechanisms. A total of 32 *Octopus mimus* adult females were sampled during ovarian maturation; biochemical composition (metabolites and digestive enzymes) of digestive gland (DG) and ovaries (only metabolites) were followed during physiological and functional maturation. Levels of protein (Prot), triacyl glycerol (TG), cholesterol (Chol), glucose (Glu) and glycogen (Gly) were evaluated. The activity of alkaline and acidic enzymes also was measured in DG. Simultaneously, groups of eggs coming from mature females were sampled along development, and metabolites (Prot, TG, Glu, Gly, TG, Chol), digestive enzymes activity (Lipases, alkaline and acidic), antioxidant defence mechanisms and radical oxygen species (ROS) were evaluated. This study shows that ovarium is a site for reserve of some nutrients for reproduction. Presumably, TG where stored at the beginning of the maturation processes followed by Chol, both at the same time were energetically supported by Glu, derived from Gly following gluconeogenic pathways. Nutrients and enzymes (metabolic, digestive and REDOX system) where maintained without significant changes and in a low activity during organogenesis. Our findings suggest that activity was not energetically costly; in contrast, during the embryo growth there was mobilization of nutrients and activation of the metabolic and digestive enzymes. Increments in consumption of yolk and glycogen, and reduction in molecules associated with oxidative stress allowed paralarvae to hatch with the antioxidant defence mechanisms ready to support ROS production.

## Introduction

Sexual maturation and reproduction influence the status of a number of physiological processes and consequently animal ecology and behaviour (Zamora & Olivares 2004). Studies on *Octopus vulgaris, O. defilippi* (Rosa et al. 2004) and on the Ommastrephid squids *Illex coindetii* and *Todaropsis eblanae* (Rosa et al. 2005) suggest that those species take the energy for egg production directly from food rather than from stored products in digestive glands or muscle, demonstrating the importance of trophic ecology for reproduction in those species. Laboratory studies have confirmed that the type of food has a strong influence in the biochemical characteristics of octopus eggs, embryos and hatchlings. The type of diet (fresh or formulated) during *O. maya* female reproduction has an effect on biochemical and morphological characteristics of embryos and hatchlings (Caamal-Monsreal et al. 2015; Tercero-Iglesias et al. 2015). *Octopus maya* females fed with mixed diets produced yolk of higher quality, allowing hatchlings a better performance during the first days of culture compared with hatchlings from females fed with mono-diet. In addition, nutrition influences the health condition of *O. maya* females and determined the capacity of the animals to maintain their physiological integrity during the post-spawning period (Caamal-Monsreal et al. 2015; Tercero-Iglesias et al. 2015), when females require enough energy to protect the spawn during the embryo development (Roumbedakis et al. 2017). A mixed maternal diet resulted in more hatchlings from *O. vulgaris* females than from females fed with mono-diet (only crabs) (Márquez et al. 2013). This finding suggests that the relationship between the nutrients of the diet (AAs and FAs), metabolic pathways and embryo characteristics previously observed in *O. maya* (Caamal-Monsreal et al., 2017) may operate similarly in other octopus species. Despite the recent advances on the importance of the type of food on embryos and hatchlings characteristics, there is no information on the biochemical dynamics associated with the processing of nutrients during the maturation of females, how nutrients are stored in eggs and used by embryos for their development. There are physiological changes during octopus embryo development associated with the use of nutrients stored in yolk to synthesize organs and tissues. Caamal-Monsreal et al., (2016) and Sánchez-García et al., (2017) showed that from stage XV onwards the yolk consumption of *O. maya* embryos was significantly higher than observed in previous stages. This suggests that embryo metabolism is accelerated at stage XV to stimulate the growth of embryos after organogenesis. From stage XV joint with energetic metabolism, there is an increment in the activity of catabolic enzymes that transform yolk in molecules that are physiologically useful for embryos (Caamal-Monsreal et al. 2016). That mobilization of reserves causes, on the one hand, an increment in oxygen consumption with consequences in energy production. On the other hand, it causes the production of radical oxygen species (ROS) due to the increment of metabolic rate (Repolho et al. 2014; Sánchez-García et al. 2017). Octopus embryos are not capable of eliminating ROS efficiently when exposed to thermal changes (Repolho et al. 2014; Sánchez-García et al. 2017). However, there is no information on the relationship between the use of yolk reserves, the antioxidant defence mechanisms, and ROS production during octopus embryos development.

*Octopus mimus* is one of the most important species of octopus in the Pacific Ocean further South of Ecuador. This species inhabits off northern Peru to San Vicente bay in Chile, where it sustains important artisanal benthic fisheries grounds in both countries (Cardoso et al. 2004; Cortez et al. 1999; Olivares et al. 1996). *Octopus mimus* reproduces throughout the year (Cardoso et al., 2004; Castro-Fuentes et al., 2002; Olivares et al., 1996). Egg laying by an individual can extend over 20 d, due to asynchrony in oocyte development and the loss of ovarian function due to their semelparous reproductive strategy (Zamora and Olivares, 2004). The time span of embryonic development changes with environmental temperature; for instance, it spans around 68 d at 16 °C during winter, whereas it lasts between 38 and 43 d at 20 °C during summer (Castro-Fuentes et al., 2002; Warnke, 1999). The optimum temperature for *O. mimus* embryos in laboratory conditions is in the range of 15 to 18°C (Uriarte et al. 2012), while one to three d old paralarvae have a thermal preference of 23 to 26°C (Zuñiga et al. 2013). There is a fair knowledge on production and rearing of *O. mimus* paralarvae in controlled conditions; for instance, environmental variables, type of tanks, light, diet (Cortez et al. 1995; Olivares et al. 1996; Zuñiga et al. 1995), histology, biochemistry and reproduction (Cortez et al. 1995; Olivares et al. 2017; Olivares et al. 2003; Olivares et al. 2001; Zamora & Olivares 2004). Recent studies have progressed on embryo development (Castro-Fuentes et al. 2002; Uriarte et al. 2012; Warnke 1999), thermal tolerance of paralarvae (Zuñiga et al. 2013), growth and nutrition (Baltazar et al. 2000; Carrasco & Guisado 2010; Gallardo et al. 2017), taxonomy and genetics (Perez-Losada et al. 2002; Söller et al. 2000). Those studies were stimulated due to the potential for aquaculture of *O. mimus* in Chile and to better understand the potential effects of environmental changes in the natural populations.

However, there is fragmented and dispersed information about the relationship between the physiological characteristics of cephalopod females, and the physiology of embryos and hatchlings. This information will allow to establish a reference for comparison of physiological process that occurs from brood stock to the next generation. In this sense, the present study is focused in biochemical composition changes that take place during gonadal maturation of octopus females and its consequences in embryo and hatchlings characteristics, putting special attention to energetic metabolites, digestive enzymes and antioxidant defence mechanisms of *O. mimus*.

## 2. Material and methods

### 2.1 Animals

A total of 32 *O. mimus* females (1179 ± 651 g BW) were collected by scuba diving using the gear hook (the local method) at 1–5 m depth off the coast of Antofagasta, Chile (23°38’39 S, 70°24’;39 W). All captured females above these sizes were anatomically ready with a developed reproductive system (Zuñiga *et al*. 2014). This study was approved by the Animal Care committee of Universidad de Antofagasta, Chile and complied with the Experimental Animal Ethics Committee of the Faculty of Chemistry at Universidad Nacional Autónoma de México (Permit Number: Oficio/FQ/CICUAL/099/15).

### 2.2 Reproductive condition

In the laboratory 23 females were immersed in cold sea water (6–8°C) to induce loss of sensation and to enable humane killing (Andrews *et al*. 2013) as suggested for sub-tropical cephalopod species (Roper and Sweeney 1983). The organisms were dissected immediately after dormancy and four weights (±0.001 g) were recorded per octopus: BW, g–total body weight; Ovw, g – ovarium weight; RSW–reproductive system weight, identified as Ovarium with oviducts and oviducal glands; and DG, g - digestive gland weight (g). Eviscerated wet weight was measured, and DG and ovarium samples were placed in Eppendorf tubes and freeze at −80°C. Reproductive system weight, gonadosomatic and digestive gland indexes were calculated as follows:

Reproductive system weight (RSWI,%) = (RSW/BW − RSW) × 100

Gonadosomatic index (GI,%) = (Ovw/BW − Ovw) × 100

Digestive gland index (DGI, %) = (DG/BW − DG) × 100

*Octopus mimus* females were classified based on their maturity status; females were immature and 15 were at gonadal maturation. Maturating females were classified in three categories: 1) physiological maturation (Phys Mat; n = 4), 2) early functional maturation (Ea Func Mat; n = 6), and 3) late functional maturation (La Func Mat; n = 5). Phys Mat animals were at initial vitelogenesis (stage III of ovocytes), Ea Func Mat were females with eggs in the reproductive coeloma and La Func Mat were females with eggs at the end of the maturation process (Avila-Poveda et al. 2016; Olivares et al. 2017). To classify maturing females, preserved (formalin, pH 7), ovarium samples (one per each sampled female, n= 23) were cut at the middle level in transverse and longitudinal sections. The sections were washed in 70% ethanol and dehydrated in ethanol series, cleared in benzene, infiltrated and embedded in paraplast ®. Serial sections were cut at a thickness of 5 μm using a Leitz 1512 manual rotary microtome, mounted on glass slides and stained using the routine Harris hematoxylin-eosin regressive method (HHE_2_; Luna 1968, Howard and Smith 1983). Alcian blue at pH 1.0 was used to contrast acidic mucopolysaccharides (Humason 1962).

### 2.3 Embryo development

Eight females were placed individually in 108 L tanks with open and aerated seawater flow at optimum temperatures of 16 to 20°C (Uriarte et al. 2012). These individuals were kept in a semi-dark environment to stimulate spawning conditions, which occurred 8 to 20 d later. A string of eggs from each female was sampled every 4 to 7 d; embryos were classified, sorted and freeze dried for biochemical analysis. Stages of embryo development for *Octopus mimus* were classified according to Naef (1928). One to two d old paralarvae were sampled and stored at −80°C. Samples of females, embryos and paralarvae were freeze dried and transported to Unidad Multidisciplinaria de Docencia e Investigación of Faculty of Science, of Universidad Nacional Autónoma de México located in Sisal, Mexico for biochemical analysis.

### 2.4 Metabolites, digestive enzymes, antioxidant mechanisms and AChE during maturation process, embryo development and hatchlings

#### Metabolites

Samples were homogenized in cold buffer Tris pH 7.4 at 100 mg tissue/mL using a Potter-Elvehjem homogenizer. The samples were then centrifuged at 10,000 × g for 5 min at 4°C and the supernatant was separated and stored at −80°C until analysis. Acylglycerols (AG) were analyzed using the Ellitech TGML5415 kit; cholesterol was measured using the Ellitech CHSL5505 kit, and glucose was analyzed using the Ellitech GPSL0507 kit following manufacturer’s instructions. Samples were diluted 1:300 for soluble protein determination using a commercial kit (Bio-Rad; Cat. 500-0006) (Bradford 1976). Determinations were adapted to a microplate using 20 μL of supernatant and 200 μL of enzyme chromogen reagent. Absorbance was recorded using a microplate reader (Benchmark Plus, Bio-Rad), concentrations were calculated from a standard substrate solution and expressed as mg/ml.

Enzyme activity assays. Acid proteases activity at pH 4 was evaluated according to Anson (1938) with adjustments, using a solution of 1% (w/v) hemoglobin (BD Bioxom- USB Products – 217500 hammersten quality) in universal buffer (Stauffer, 1989). Alkaline proteases activity of the extracts at pH 8 was measured using the method of Kunitz (1947) modified by Walter (1984), using 1% (w/v) casein (Affymetrix, 1082-C) as substrate in 100 mM universal buffer. In both assays, 0.5 ml of the substrate solution was mixed in a reaction tube with 0.5 ml of universal buffer and 20 μl of enzyme preparation (1:100 dilution) and incubated for 10 min at 35 °C and 40 °C for alkaline proteases and acid proteases, respectively. The reaction was stopped by adding 0.5 ml of 20% (w/v) trichloroacetic acid (TCA) and cooling on ice for 15 min. The precipitated undigested substrate was separated by centrifugation for 15 min at 13,170 g. The absorbance of the supernatants was measured spectrophotometrically at 280 nm. Control assays (blanks) received a TCA solution before the substrate was added.

For trypsin, the adjusted methods of Charney and Tomarelli (1947) was used. The trypsin activity of the extracts was measured using 1 mM BAPNA (Benzoyl-L-Arg-*p*-nitroanilide) as substrate TRIS 0.1 M at pH 7. This assay was made in microplate as follows: 250 μl of the substrate solution was mixed with 5 μl of enzyme preparation (1:2). The absorbance was measured spectrophotometrically at 405 nm every minute, during 2 minutes. A mix of 5 μl of distilled water with 250 μl of substrate solution was used as control assay (blank).

Lipases activity was measured in microplate using 200 μl substrate solution (TRIS 0.5 M, pH 7.4, sodium taurocholate 5 mM, sodium chloride 100 mM, 4 nitrophenyl octanoate 0.35 mM) and 5 μl homogenate diluited (1:2) in TRIS 0.5 M, pH 7.4 (Gjellesvik, et al. 1992). Absorbance was read at 415 nm every minute, during 10 minutes.

For all enzymes, units of activity were expressed as change in absorbance per minute per milligram of protein (U=Abs_nm_ min^−1^mg^−1^ protein).

Antioxidant defence system. Samples were snap-frozen in liquid nitrogen, lyophilized and stored at −20°C until homogenization. Samples were homogenized in cold buffer Tris 0.05 M pH 7.4 at 10 mg tissue/mL using a Potter-Elvejhem homogenizer. For enzyme activity assays, homogenates were centrifuged at 10,000 × g for 5 min at 4°C and the supernatant was separated for analysis. All samples were stored at −80°C until analysis.

Redox potential was measured with a probe (ArrowDox Measurement System, ORP-146CXS, Los Angeles, USA) in each homogenate (in mV). Posteriorly, the homogenate was divided for triplicate assays to measure the activities of acetylcholinesterase (AChE), carboxylesterase (CbE), catalase (CAT), glutathione S-transferase (GST), and for levels of lipid peroxidation (LPO) and total glutathione (GSH). All spectrophotometric measurements were realized in a micro-plate reader. AChE activity was measured using a modification of the method described by Ellman et al. (1961), which was adapted to a microplate reader (Rodríguez-Fuentes et al. 2008). Each well contained 10 μL of the enzyme supernatant and 180 μL of 5, 5’ -dithiobis (2 nitrobenzoic acid) (DTNB) 0.5 M in 0.05 mM Tris buffer pH 7.4. The reaction started by adding 10 μL of acetylthiocholine iodide (final concentration 1 mM) and the rate of change in the absorbance at 405 nm was measured for 120s. CbE activity, a detox enzyme, was measured using ρ-nitrophenyl-α-arabinofuranoside (ρNPA) substrate, as indicated by Hosokawa and Satoh (2001) with some modifications (25 μL of the supernatant and 200 μL of ρNPA were mixed, and the reaction was recorded for 5 min at 405 nm).

CAT activity was measured using the Goth (1991) method with modifications made by Hadwan and Abed (2016). In this method, the undecomposed H_2_O_2_ is measured after 3 minutes with ammonium molybdate to produce a yellowish color that has a maximun absorbance at 374 nm. GST activity was determined from the reaction between reduced glutathione and 1-chloro-2.4-dinitrobenzene at 340 nm (Habig and Jakoby 1981). AChE, CbE, CAT, and GST activities were reported in nmol min^−1^ mg protein^−1^. Proteins were analyzed in the supernatant according to Bradford (1976) and was used to normalize enzyme activities. Total glutathione (GSH) was measured with a Sigma-Aldrich Glutathione Assay Kit (CS0260). This kit utilizes an enzymatic recycling method with glutathione reductase (Baker *et al*., 1990). The sulfhydryl group of GSH reacts with Ellman’s reagent and produces a yellow colored compound that is read at 405 nm. LPO was evaluated using the PeroxiDetect Kit (PD1, Sigma-Aldrich, USA) following the manufacturer’s instructions and was reported in nmol mL^−1^. The procedure is based on peroxides oxide iron (Fe^3+^) that forms a coloring component with xylenol orange at acidic pH measured at 560 nm.

### 2.5 Statistical analysis

General data were expressed as mean (± standard deviation, SD). Differences of the mean values of each measurement variables, female tissue, embryo and paralarvae (i.e. BW, Ovw, RSW, and DG weights) were tested each one using one-way ANOVA followed by Fisher LSD (least significant difference) test (Sokal and Rohlf 1995). Before ANOVA, assumption tests were carried out to determine the homogeneity of variances for each measurement and those that did not fit the premises for ANOVA were transformed using more appropriate measurement scales (McCune *et al*. 2002). Statistical analyses were carried out using STATISTICA^®^ 7. The significance of the statistical difference was accepted at p < 0.05.

We evaluated the synergistic way in which the physiological processes are carried out in *O. mimus* females during the maturation of the gonads and in embryos during development. Therefore, multivariate sets of descriptors were analysed: 1) reproductive condition: BW, OVW, DG, EBW and RSW (mg); 2) metabolite concentration in ovaries: Glycogen, Glucose, Chol, Triacyl and Prot (mg/ml); 3) metabolite concentration and enzyme activity in the digestive gland: Glycogen, Glucose, Chol, Triacyl and Prot, AcidProt and AlkaProt (UI/mg protein). Data on *O. mimus* embryos and paralarvae were analysed using three multivariate sets of descriptors considering four well recognized phases during embryo development: i) organogenesis (namely Pre); ii) the end of organogenesis around stages XIV and XV (namely Organ); iii) Growth (post organogenesis, namely Post) and iv) one d old (1^st^) and two d old (2^nd^) paralarvae. The descriptors used were 1) metabolite concentration in body as a whole: Glycogen, Glucose, Chol, Triacyl and Prot (mg/ml); 2) enzyme activity: acid and alkaline proteases (UI/mg protein) and 3) antioxidant defence mechanisms: Ache, CbE, GsT, SOD, CAT (nMol/min/mg protein), ORP (mV), GsH (nMol/ml), LPO (nMol peroxide/ml). Principal Coordinate Analysis (PCoA) was applied on Euclidean distance matrices of samples in each data set (Legendre and Legendre, 1998). Data were square root (female data) or log-transformed (embryo and paralarvae data), and normalised by centring and dividing between the standard deviation of each variable prior to analysis (Legendre and Legendre, 1998).

A permutational multiple ANOVA was applied on the distance matrices to detect differences amongst female octopuses in four different stages of gonadic maturation (fixed factor: Imm, PhyMat, EarFunMat, LatFunMat), and amongst embryos and paralarvae in different stage of development (Pre, Organ, Post, 1st and 2nd paralarvae). Permutational multiple paired t-tests were used to compare the centroids of the different stages in each data set; 9999 unrestricted permutations of raw data were used to generate the empirical *F* and *t*-distributions (Anderson, 2001; McArdle and Anderson, 2001).

## 3. Results

### 3.1 *Octopus mimus* females reproductive condition

Total and eviscerated wet weight changed with maturation of females with low values in females at immature stage and high values in females in late functional maturity stage (Fig. 1A). At the end of maturation, total and eviscerated weight resulted 3.2 and 2.7 higher than observed in immature females (p < 0.0001; Fig. 1A). Likewise, an increment of reproductive complex system (RCS), ovarium and oviductal wet weight were registered along the maturity stages; RSW, Ovw and oviductal gland wet weight values at the end of maturity stages were 108, 135 and 2.55 times higher than in immature females, respectively (Fig 1B). As a consequence, increments in RSWI and OvwI were observed along the maturity stages (Fig. 1C).

**Fig. 1.**
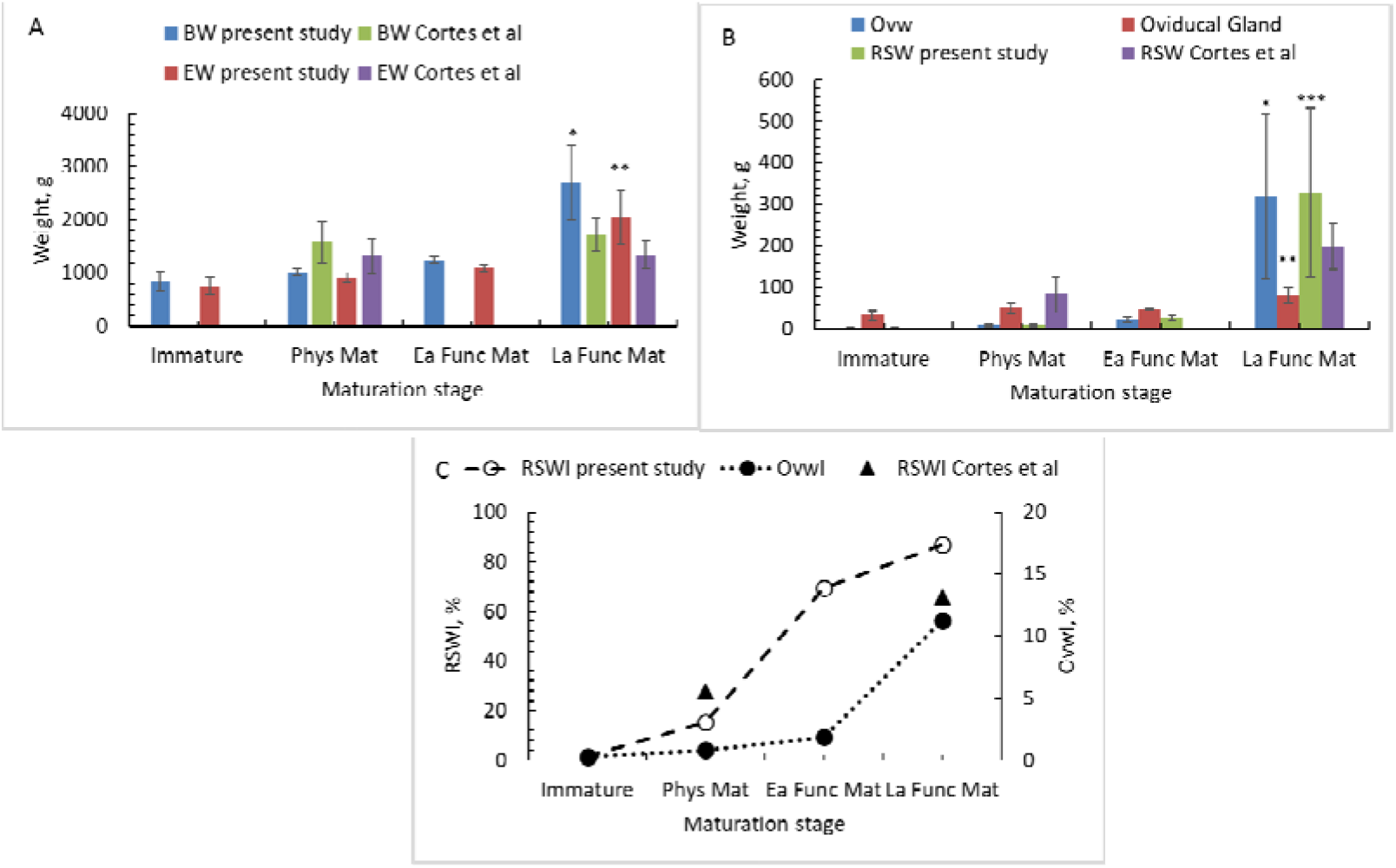
Total, eviscerated (A), reproductive tract (B), and gonadosomatic and reproductive system indices (C) of *O. mimus* females along the maturation stages obtained in the present study and by Cortes et al. (1995). Physiological maturity (Phys Mat) was obtained from females with a OvwI < 2 and in stage III of oocytes development. Early functional maturation stage (Ea Fun Mat), was recognized from females with eggs in the reproductive coelom and with OvwI > 2. Late functional maturation stage (La Func Mat), was obtained from females close to spawn (Arkhipkin, 1992; Avila-Poveda *et al*., 2016; Olivares *et al*., 2017). RSWI = reproductive system index; OvwI = ovarium Index.

Digestive gland wet weight changed along the maturity stages with low values in immature females and high values at the end of the maturation process (p < 0.001; Fig. 2). In contrast, hepatosomatic index (HI, %) increased with maturation stages to reach a peak at the beginning of the functional maturation, when oocytes were send to reproductive coelom (Fig. 2). The hepatosomatic index was reduced at the end of the maturation stage.

**Fig. 2.**
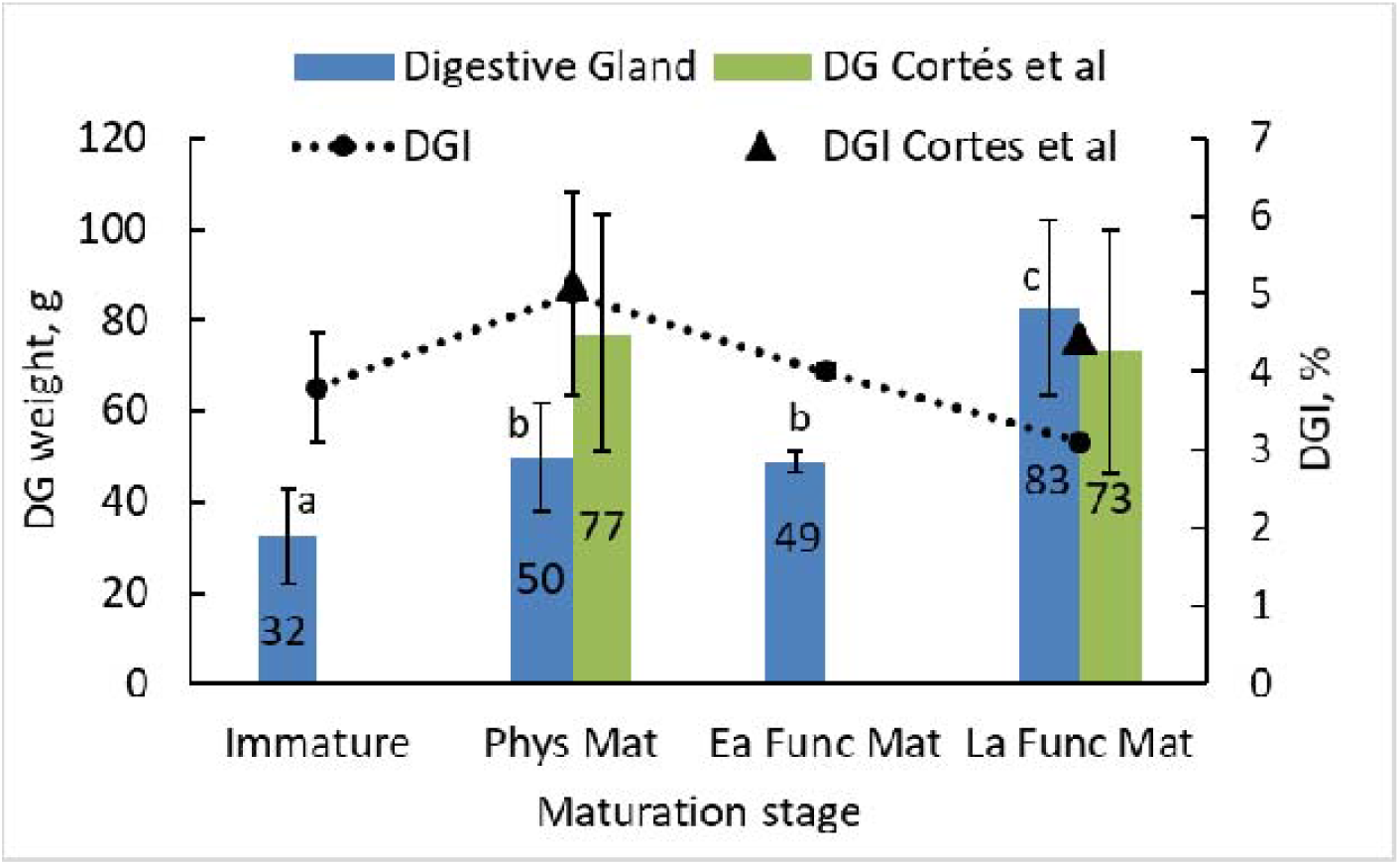
Digestive gland weight (DG, g) and digestive gland index (DGI, %) of *O. mimus* females along the maturation stages obtained in the present study and by Cortes et al. (1995). Physiological maturity (Phys Mat) was obtained from females with a IGS < 2 and in stage III of oocytes development. Early functional maturation stage (Ea Fun Mat), was recognized from females with eggs in the reproductive coelom and with OvwI > 2. Late functional maturation stage (La Func Mat), was obtained from females close to spawn (Arkhipkin, 1992; Avila-Poveda, *et al*., 2016; Olivares, *et al*., 2017)

### 3.1 Biochemical composition

#### a. Glycogen

Immature *O. mimus* females showed digestive gland glycogen levels 59% lower than females at the beginning of the maturation process (Fig. 1A; p < 0.001). A significant increment of DG glycogen was subsequently registered in females in physiological maturation, reaching values 2.4 times higher than the levels observed in immature females (p < 0.001; Fig. 3A). A reduction in 38% of digestive gland glycogen values was registered (p < 0.01; Fig. 3A). Increments of ovarium glycogen concentration were detected during maturation with high glycogen levels obtained in the ovaries of females at the end of the maturation process (Fig. 3A; p < 0.01). In *O. mimus* embryos, glycogen levels of stages I to XIV and XVIII did not show statistical differences with a mean value of 26.8 ± 9.0 mg/g (p > 0.05; Fig. 3A). Glycogen levels of embryos at stage XVI and paralarvae 1 and 2 were statistically lower than the rest of the embryos (Fig. 3A; p < 0.01).

**Fig. 3.**
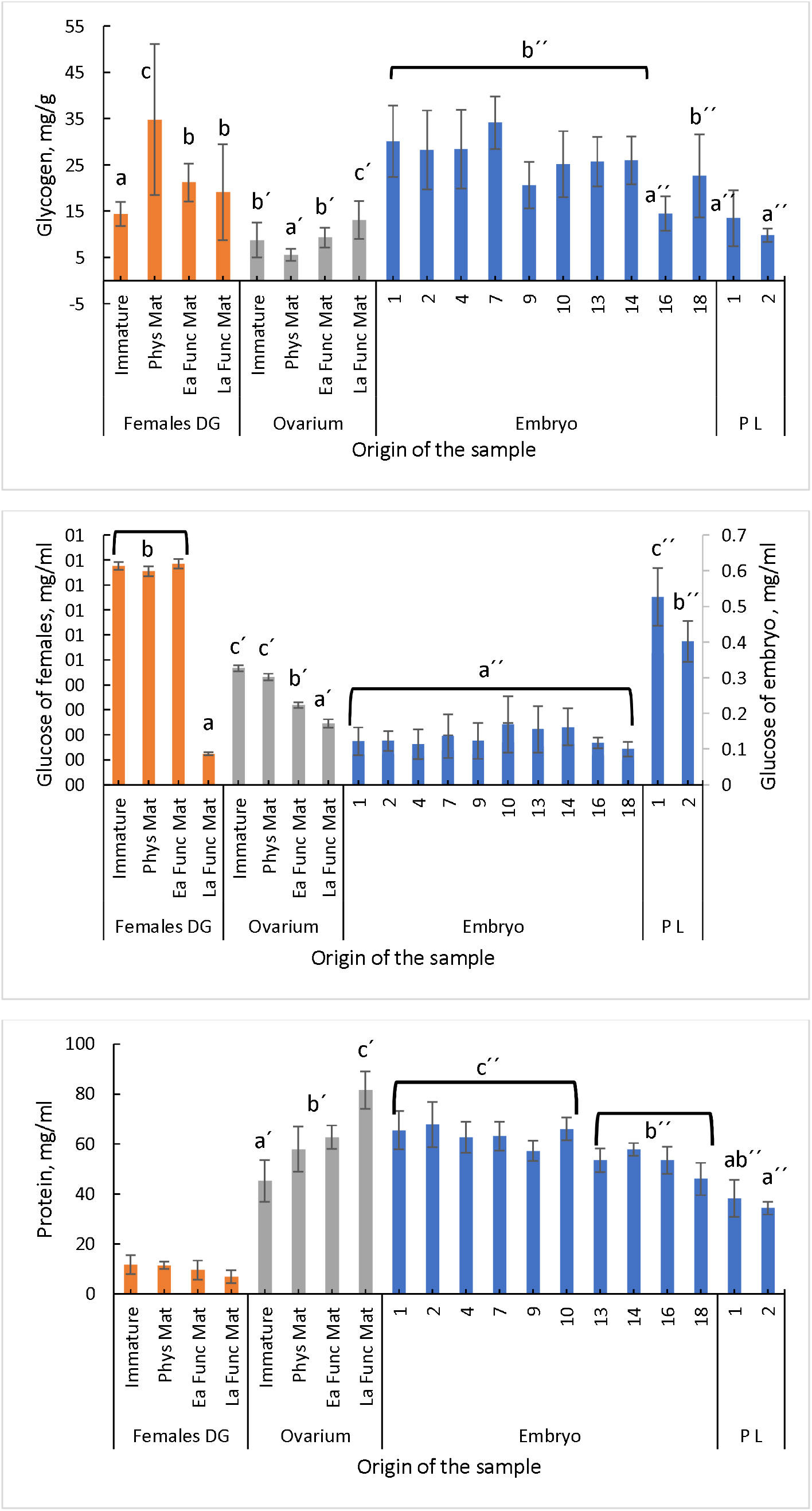
Glycogen, glucose and protein of *O. mimus* females (digestive gland: DG) and ovarium), embryos and paralarvae (PL) maintained in laboratory conditions at 16°C. Values as mean + SD. Different letters indicate statistical differences at P < 0. 05 level.

#### b. Glucose

Digestive gland glucose levels were similar in immature females and during almost all maturation process (p > 0.5; Fig 3B). Only females in late functional maturation process had glucose levels significantly lower than in the rest of the maturation conditions (p < 0.001; Fig. 3B). A reduction in glucose levels was recorded in ovaries according with the maturation process, with higher levels in immature females (0.5 mg/ml) than in late functional maturation condition (p < 0.001; Fig 3B). Glucose levels were similar along embro development. In consequence, a mean value of 0.13 ±0.02 mg/ml was calculated for all embryo stages (p > 0.05; Fig 3B). Glucose levels in 1 d old paralarvae were 4 times higher than in embryos and 1.3 times higher than observed in 2 d old paralarvae (p < 0.002; Fig 3B).

#### c. Protein

Digestive gland protein did not change during the female maturation process (p > 0.05; Fig. 3C). In contrast, an increment of soluble protein was recorded in ovarium in relation with maturation process, with low values in immature females and high at the end of the maturation process (p < 0.001; Fig. 3C). There were no statistical differences in protein levels of embryos of stages I to X, with a mean value of 63.8 ± 3.7 mg/ml (p > 0.05; Fig. 3C). After those stages, a reduction of soluble protein levels occurred in embryos and paralarvae to reach the lower value in 2 d paralarvae of *O. mimus* (p > 0.05 Fig; 3B).

#### d. Triacyl glycerol (TG)

Digestive gland TG values did not change with female maturation stages; for this reason, a mean value of 0.35 ± 0.08 mg/ml was calculated (p > 0.05; Fig. 4A). There were strong changes in TG values in female ovarium, with low values in immature females (1.05 mg/ml) and at the end of the maturation stage (1.6 mg/ml) (p > 0.05). In contrast, TG values were high in females at the end of physiological maturation and during the early maturation process (p < 0.001; Fig. 4A). Levels of TG in embryos were highly variable; TG levels were high in stages I, IV, VII, and X to XVIII but low in stages II and IX (p < 0.01; Fig. 4A). Values of TG recorded in 1 and 2 d old paralarvae resulted 1.5 times higher than the maximum value recorded in embryos at stage X (p < 0.01; Fig. 4A).

**Fig. 4.**
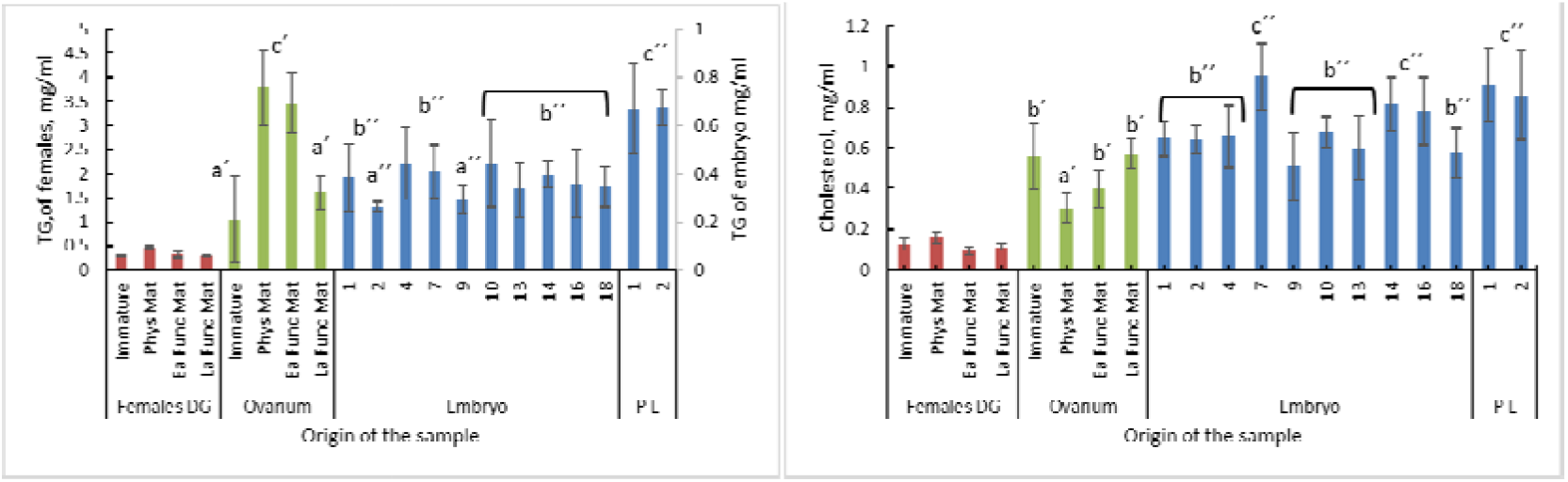
Triacyl glycerol and cholesterol of *O. mimus* females (digestive gland: DG) and ovarium), embryos and paralarvae (PL) maintained in laboratory conditions at 16°C. Values as mean + SD. Different letters indicate statistical differences at P < 0. 05 level.

#### e. Cholesterol

Cholesterol levels in the digestive gland remained constant along the maturation stages of *O. mimus* females (p > 0.05; Fig. 4B). Cholesterol levels in the ovarium changed with maturation stages of females with high levels in immature animals. There was a subsequent reduction of cholesterol levels at the beginning of the maturation process, followed by an increment in the next maturation stages to reach the maximum concentration level at the end of the maturation process (p < 0.002; Fig. 4B).

### 3.2 Digestive enzymes

Acidic and alkaline proteases activity was constant along the embryo development until stage XIV by alkaline and stage XVIII by acidic proteases (Fig. 5). Alkaline proteases activity increased after stage XIV to reach its maximum and significant activity after hatching (p < 0.001; Fig 5A). A maximum acidic proteases activity also was recorded in *O. mimus* after hatching. In adult females, there were no statistical differences in activity of acidic proteases along the maturity stages (p > 0.05; Fig 5A). In contrast, a lower activity of alkaline proteases was registered in immature females in comparison with maturating and mature *O. mimus* females (Fig. 5A). Lipases and trypsin activities changed with embryo development, with low values in stages lower than XIII, and higher in stages XIV to XVIII (p < 0.001; Fig. 5B). A reduction on lipases activity was recorded in 1 d old paralarvae (p < 0.001; Fig. 5B).

**Fig. 5.**
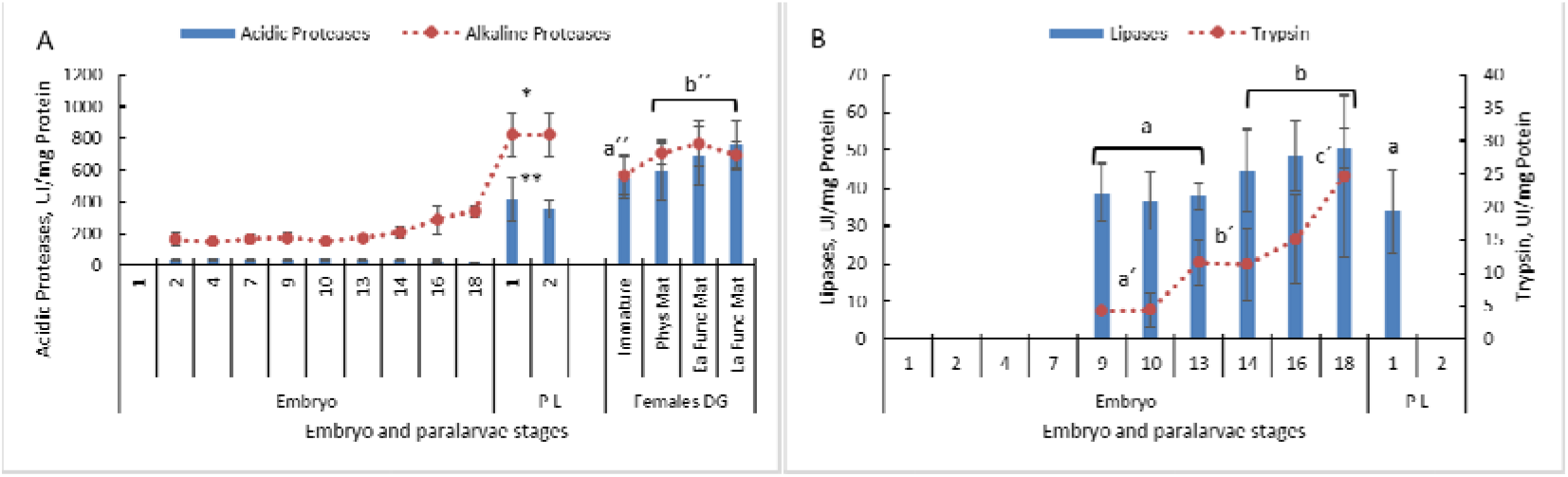
Digestive enzymes activity in entire embryos, paralarvae (PL) and in the digestive gland of *O. mimus* females obtained in animals maintained in laboratory conditions at 16°C. Values as mean + SD. Different letters or asterisk indicate statistical differences at P < 0.05 level.

3.3. Acetyl cholinesterase (AChE) and antioxidant defence mechanisms (ADM) AChE increased along the embryo development with low AChE values at the beginning of embryo development and high values after hatching (p < 0.01; Fig 6A). Antioxidant defence mechanisms (GST, SOD and CAT) showed the same tendency, with low values at the beginning of embryo development and high values after hatching (p < 0.01; Fig. 6 B to D). As a consequence, a reduction in oxidative damage (CbE; ORP and LPO) and GSH was observed with high values in embryos at the beginning of the development and low values after hatching. It is interesting to note that LPO and GSH levels start to be reduced around stage XV to maintain a low level until hatch (p < 0.001; Fig 6).

**Fig. 6.**
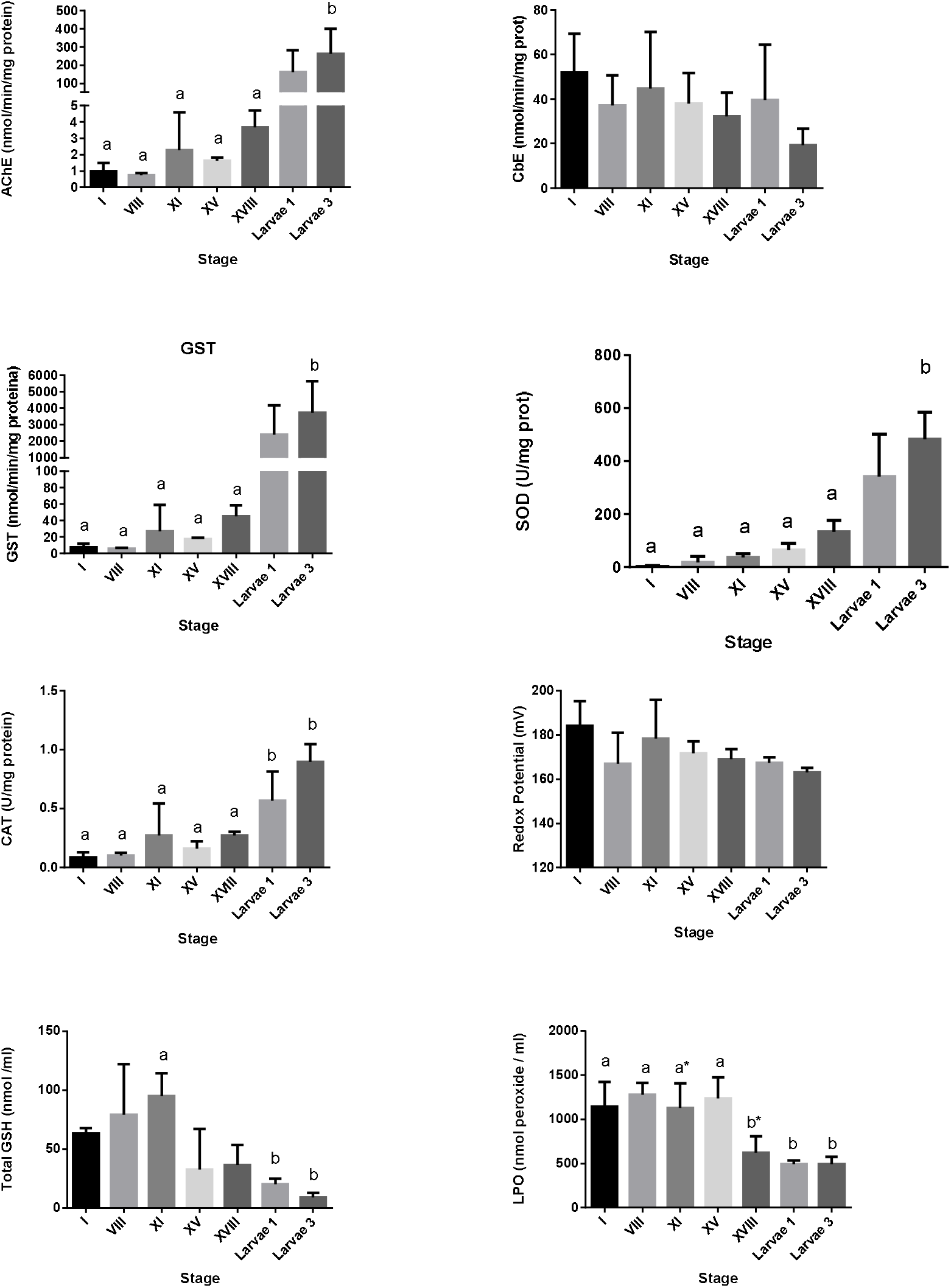
Antioxidant defence mechanisms and acetyl cholinesterase activity of embryos and paralarvae of *O. mimus* obtained in laboratory conditions at 16°C. Values as mean + SD. Different letters means statistical differences between treatments at p < 0.05 level.

Ordination by PCoA of the reproductive condition of female *O. mimus* showed that the PCo1 explained 94.9 % of total variation in the data, with EBW, BW, OVW and RSW greatly contributing to ordination in the horizontal axis (Fig. 7A). The PCo2 explained only 4.2% of the total variation, with differences in DG contributing to the ordination in the vertical axis. Values of all the reproductive condition descriptors increased as females advanced from immature to physiological and functional stages of gonadic maturation (Fig. 7A). Females in late functional maturity were distinctly separated and associated WITH the highest weight values.

**Fig. 7.**
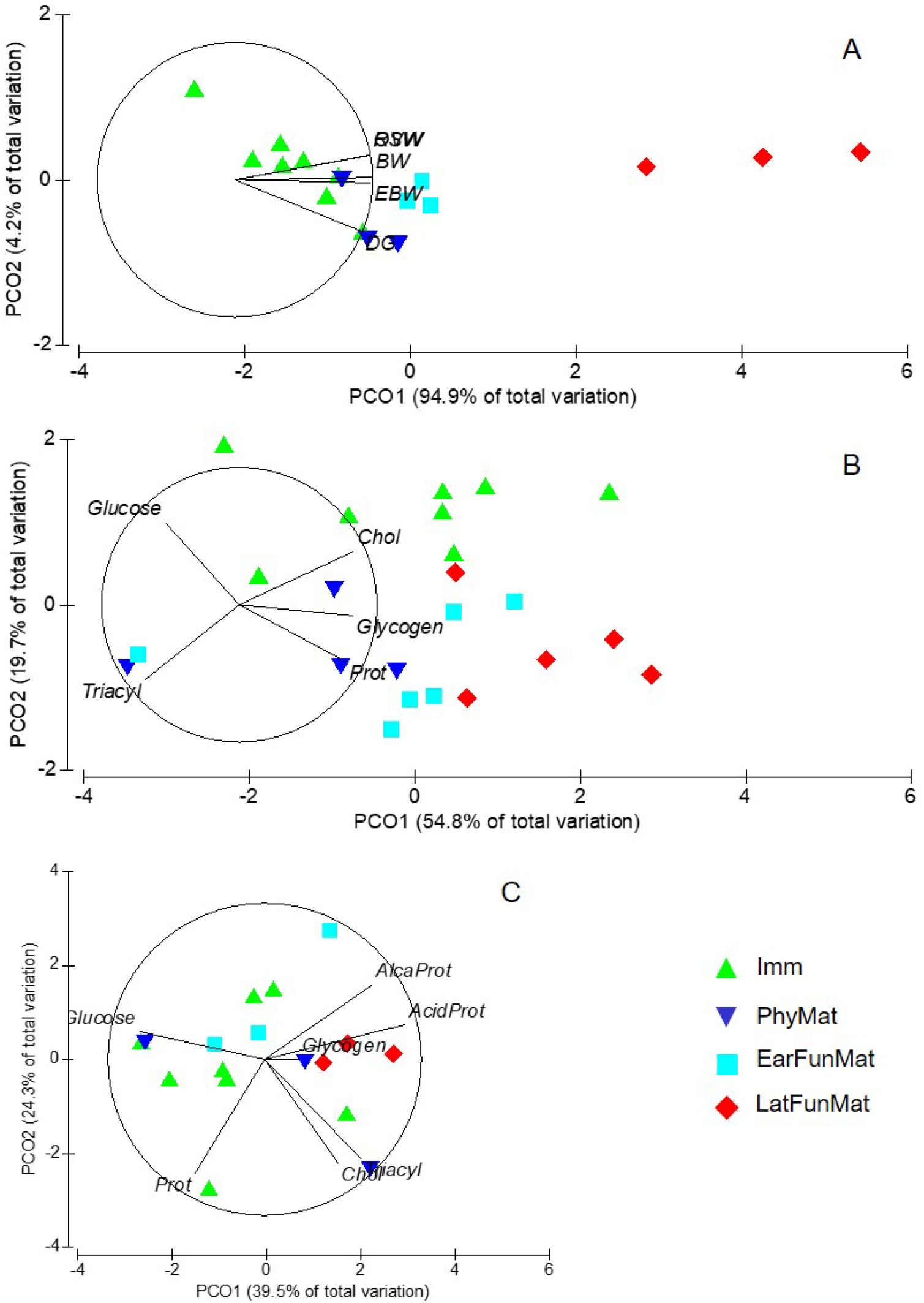
Principal coordinate analysis (PCO) of females of *O. mimus* during ovarian maturation. A. Body weight characteristics (Total body weight: BW; Eviscerated weight: EBW; reproductive tract weight: RTW; Digestive gland weight: DG). B. ovarian metabolites (Chol: Cholesterol; Prot: protein; Triacyl: triacyl glycerol). C. Digestive gland metabolites and digestive enzymes (Chol: Cholesterol; Prot: protein; Triacyl: triacyl glycerol; Alcal prot: Alkaline proteases; Acid Prot: Acidic Proteases). Measures made in immature (imm), physiological maturity (PhyMat), early functional maturity (EarFunMat) and late functional maturity (LatFunMat).

The multiple ANOVA showed overall significant differences between stages of gonadic maturation (Table 1). However, paired comparisons amongst centroids revealed that only immature females and those in either early or late functional maturity differed significantly in the reproductive condition descriptors measured (Table 2).

**Table 1.**
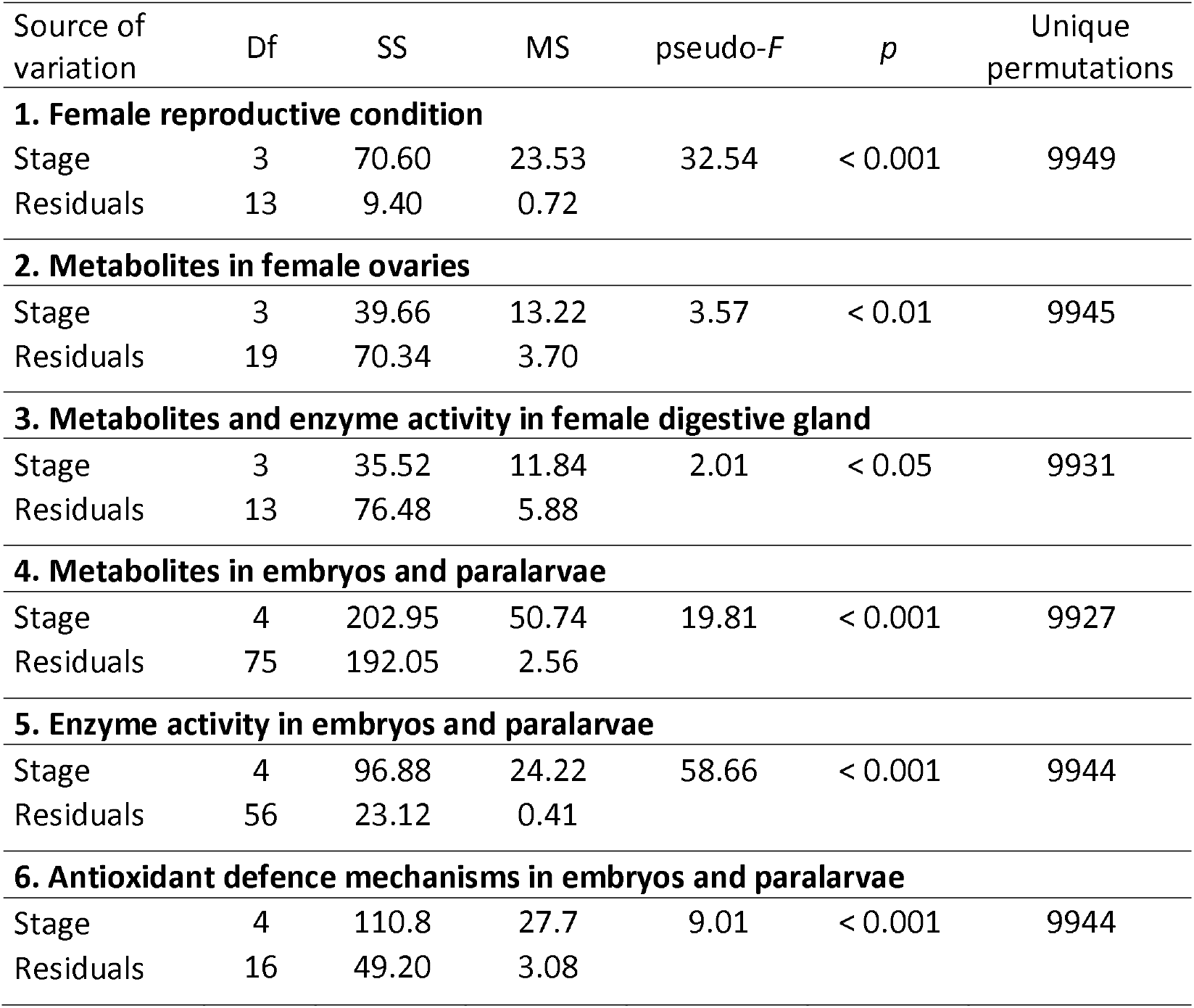
Results of one-way permutational multiple MANOVAs applied on six multivariate sets of data obtained from *O. mimus*: 1) female reproductive condition; 2) metabolite concentration in female ovaries; 3) metabolite concentration and enzyme activity in the digestive gland of female; 4) metabolite concentration; 5) enzyme activity; and 6) antioxidant defence mechanisms in embryos and paralarvae. The degrees of freedom (df), multivariate sum of squares (SS), mean square (MS), pseudo-F and p-values, and the number of unique permutations is given for each test.

| Source of variation | Df | SS | MS | pseudo-*F* | *p* | Unique permutations |
| --- | --- | --- | --- | --- | --- | --- |
| **1. Female reproductive condition** |  |  |  |  |  |  |
| Stage | 3 | 70.60 | 23.53 | 32.54 | < 0.001 | 9949 |
| Residuals | 13 | 9.40 | 0.72 |  |  |  |
| **2. Metabolites in female ovaries** |  |  |  |  |  |  |
| Stage | 3 | 39.66 | 13.22 | 3.57 | < 0.01 | 9945 |
| Residuals | 19 | 70.34 | 3.70 |  |  |  |
| **3. Metabolites and enzyme activity in female digestive gland** |  |  |  |  |  |  |
| Stage | 3 | 35.52 | 11.84 | 2.01 | < 0.05 | 9931 |
| Residuals | 13 | 76.48 | 5.88 |  |  |  |
| **4. Metabolites in embryos and paralarvae** |  |  |  |  |  |  |
| Stage | 4 | 202.95 | 50.74 | 19.81 | < 0.001 | 9927 |
| Residuals | 75 | 192.05 | 2.56 |  |  |  |
| **5. Enzyme activity in embryos and paralarvae** |  |  |  |  |  |  |
| Stage | 4 | 96.88 | 24.22 | 58.66 | < 0.001 | 9944 |
| Residuals | 56 | 23.12 | 0.41 |  |  |  |
| **6. Antioxidant defence mechanisms in embryos and paralarvae** |  |  |  |  |  |  |
| Stage | 4 | 110.8 | 27.7 | 9.01 | < 0.001 | 9944 |
| Residuals | 16 | 49.20 | 3.08 |  |  |  |

**Table 2.** Results of permutational paired t-tests that compared centroids representing data in six multivariate sets of data obtained from female *O. mimus* in different stages of gonadic maturation (Imm: immature, PhyMat: physiological maturity, EarFuncMat: early functional maturity, and LatFuncMat: late functional maturity), and from embryos and paralarvae in different stage of development (Pre, Organ and Post: stages before, during, and after organogenesis, respectively, and 1st and 2nd paralarvae). Values are permutational p-values for each test.

| **1. Female reproductive condition** |  |  |  |  |
| --- | --- | --- | --- | --- |
|  | Imm | PhyMat | EarFuncMat |  |
| PhyMat | 0.053 | - | - |  |
| EarFuncMat | < 0.05 | 0.102 | - |  |
| LatFuncMat | < 0.01 | 0.100 | 0.100 |  |
| **2. Metabolites in female ovaries** |  |  |  |  |
|  | Imm | PhyMat | EarFuncMat |  |
| PhyMat | < 0.05 | - | - |  |
| EarFuncMat | < 0.05 | 0.351 | - |  |
| LatFuncMat | < 0.01 | < 0.05 | < 0.05 |  |
| **3. Metabolites in female digestive gland** |  |  |  |  |
|  | Imm | PhyMat | EarFuncMat |  |
| PhyMat | 0.225 | - | - |  |
| EarFuncMat | 0.271 | 0.498 | - |  |
| LatFuncMat | < 0.01 | 0.204 | 0.104 |  |
| **4. Metabolites in embryos and paralarvae** |  |  |  |  |
|  | Pre | Organ | Post | 1st |
| Organ | < 0.01 | - | - | - |
| Post | < 0.001 | 0.161 | - | - |
| 1st | < 0.001 | < 0.001 | < 0.001 | - |
| 2nd | < 0.001 | < 0.01 | < 0.01 | 0.567 |
| **5. Enzyme activity in embryos and paralarvae** |  |  |  |  |
|  | Pre | Organ | Post | 1st |
| Organ | < 0.001 | - | - | - |
| Post | < 0.001 | < 0.05 | - | - |
| 1st | < 0.001 | < 0.001 | < 0.01 | - |
| 2nd | < 0.001 | < 0.01 | < 0.01 | 0.617 |
| **6. Antioxidant mechanisms in embryos and paralarvae** |  |  |  |  |
|  | Pre | Organ | Post | 1st |
| Organ | 0.125 | - | - | - |
| Post | < 0.01 | 0.098 | - | - |
| 1st | < 0.01 | 0.097 | 0.098 | - |
| 2nd | < 0.01 | 0.099 | 0.096 | 0.299 |

Ordination of the concentration of metabolites in female ovaries showed that the PCo1 and PCo2 explained 74% of the total variation (54.8% and 19.2%, respectively; Fig. 7B). Concentration of Glycogen, Chol and Prot contributed mostly to the ordination on the horizontal axis, whereas Glucose and Triacyl did so on the vertical axis. The multiple ANOVA showed overall significant differences between stages of gonadic maturations (Table 1). Paired comparison between centroids allowed for three distinct groups to be formed based on ovary metabolites: immature; physiologically and early functionally mature, and late functionally mature females (Table 2).

Ovary samples from immature females were high in glucose and cholesterol, but low in triacylglicerides and proteins when compared to samples from late functionally mature females; physiologically and early functional mature females presented intermediate ovary concentrations of these metabolites (Fig. 7B).

Together, the first and second PCo only explained 63.8% of the total variation in metabolite concentration and enzyme activity of the digestive gland in female *O. mimus* (39.5% and 24.3%, respectively; Fig. 7C); the percentage of total variation increased to 79% when the third PCo was considered. Glucose was inversely correlated with glycogen, triacylglicerides and cholesterol, whilst protease activity was correlated with protein concentration in all three principal coordinates. Significant differences between stages of gonadic maturation were also detected by the general multiple MANOVA (Table 1). However, paired comparisons between centroids only revealed significant differences in immature females and those in late functional maturity (Table 2).

Immature females had high concentrations of glucose and both acid and alkaline protease activity in the digestive gland. Fully functionally mature females had relatively higher concentration of glycogen and lower protease activity in the digestive gland.

The first and second PCo explained 56.7% and 20% of total variation of metabolite concentration in embryos and paralarvae (Fig. 8A). Glucose and triacylglicerides were inversely correlated with glycogen and proteins and contributed largely to sample ordination in the horizontal axis. Cholesterol concentration contributed to order samples on the vertical axis (Fig. 8A). The multiple MANOVA detected overall significant differences amongst stages of embryo and paralarvae (Table 1). Significant differences in paired comparisons allowed to distinguish three groups: embryos in stages prior to organogenesis (Pre); embryos in stages characterised by organogenesis and immediately after (Organ and Post); and the first and second paralarvae. Embryos prior to enter organogenesis had high glycogen and protein concentration but low glucose and triacylgliceride concentration. Embryos at stages Organ and Post had high cholesterol concentrations and intermediate values in all other metabolites. The 1st and 2nd stage paralarvae had high glucose and triacylgliceride but low glycogen and protein concentrations (Fig. 8A).

**Fig. 8.**
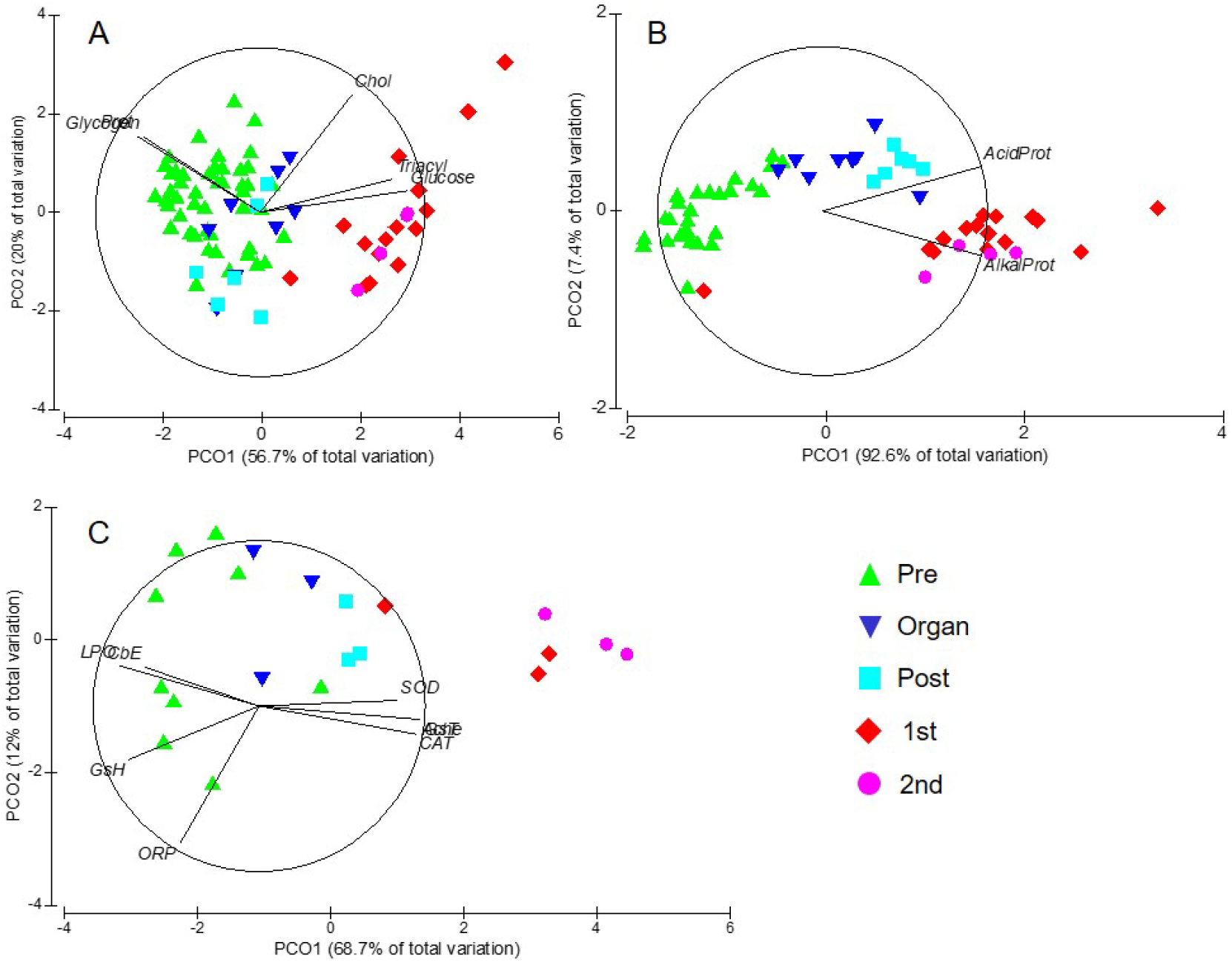
Principal coordinate analysis (PCO) of *O. mimus* embryo metabolites (A), digestive enzymes (B) and antioxidant defence mechanisms (C) measured along development. Pre = Organogenesis: stages I to XII; Organ = The end of organogenesis: stages XIV to XVI; Post = Post organogenesis: stages XVII to XIX; 1 d old Paralarvae: 1^st^; 2 d old paralarvae: 2^nd^.

Ordination of protease activity in *O. mimus* embryos and paralarvae showed that the PCo1 and PCo2 explained 92.6 and 7.4% of total variation, respectively (Fig. 8B). Enzyme activity in general increased as embryos advanced from stages before organogenesis and towards the 2nd paralarvae, with paralarvae having higher alkaline protease but lower acid protease activity than embryos. The multiple ANOVA showed significant differences between stages of development (Table 1), and significant differences were detected between all pairs of centroids except those representing the 1st and 2nd paralarvae (Table 2). These results show four distinct groups of samples regarding enzyme activity: embryos before, during and after organogenesis, and paralarvae (Fig. 8B).

Analysis of the antioxidant defence mechanisms in *O. mimus* embryos and larvae showed that the first and second PCo explained 68.7% and 12% of total data variation (Fig. 8C). Lipid peroxidation and carboxylesterase were high amongst embryos in stages before organogenesis, and were inversely correlated to acetylcholinesterase, catalase, glutathione S-transferase and superoxide dismutase, which had the highest values amongst 1^st^ and 2^nd^ day paralarvae (Fig. 8C). Whilst the multiple ANOVA showed overall significant differences throughout development (Table 1), paired tests amongst centroids revealed significant differences only between extreme stages (Table 2).

## Discussion

Previous studies of *O. mimus* from Northern Chile have investigated size at maturity and support our finding that females heavier than 2000 g are mature (Cortez et al. 1995). Patterns of body, reproductive tract, digestive gland weights and their indices observed in this study were consistent with (Cortez et al. 1995). As was expected, increments on RSW and reproductive indices were observed along the reproductive maturity stages, indicating that during ovarian development there was mobilization of energy and nutrients to reproductive organs. Studies suggested that *O. vulgaris*, *O. defilippi* (Rosa et al. 2004) and in the squids *Illex coindetii* and *Todaropsis eblanae* (Rosa et al. 2005) take the energy for egg production directly from food rather than from stored products in a specific tissue. This was concluded based on the observation that, while mature females experience increments of protein, lipid and glycogen contents in gonads, the digestive gland and muscle where without apparent changes between both maturation stages. This indicates that storage reserves were not transferred from tissue to tissue during ovaria maturation, which is consistent with our findings. Moreover, our study suggests that during the maturation process there was mobilization of nutrients at the DG and ovarium that where not observed previously. As can be expected in the DG of *O. mimus* females, free glucose is an important source of metabolic energy. Free glucose was highly concentrated along the maturation process until the early functional maturation, when oogonia are growing in the ovarium. It was observed that levels of progesterone, the hormone involved in oocytes vitellogenesis, were high during the early functional maturation of *O. maya* (Avila-Poveda et al. 2015). This finding suggests that these processes require high levels of metabolizable energy. Accordingly, glycogen and glucose in the digestive gland of *O. mimus* and *O. maya*, as well as soluble proteins were used as a source of metabolic energy during digestive processes (Gallardo et al. 2017) and growth of juveniles and pre-adults of *O. maya* (Aguila et al. 2007; Rosas et al. 2011). Hence the importance of those nutrients in the physiology of cephalopods. According to Martínez et al., (2014) this energy is the result of gluconeogenic pathways supported by the protein metabolism due to the carnivorous habits of cephalopods. For this reason we hypothesise that, as in DG (Martinez et al., 2014) in the ovarium part of the glycogen and glucose registered followed the glycogenic pathways (Hochachka & Fields 1982) (Fig. 9),as was previously described in muscle of different cephalopod species (Gallardo et al. 2017; Hochachka & Fields 1982; Morales et al. 2017).

**Fig. 9.**
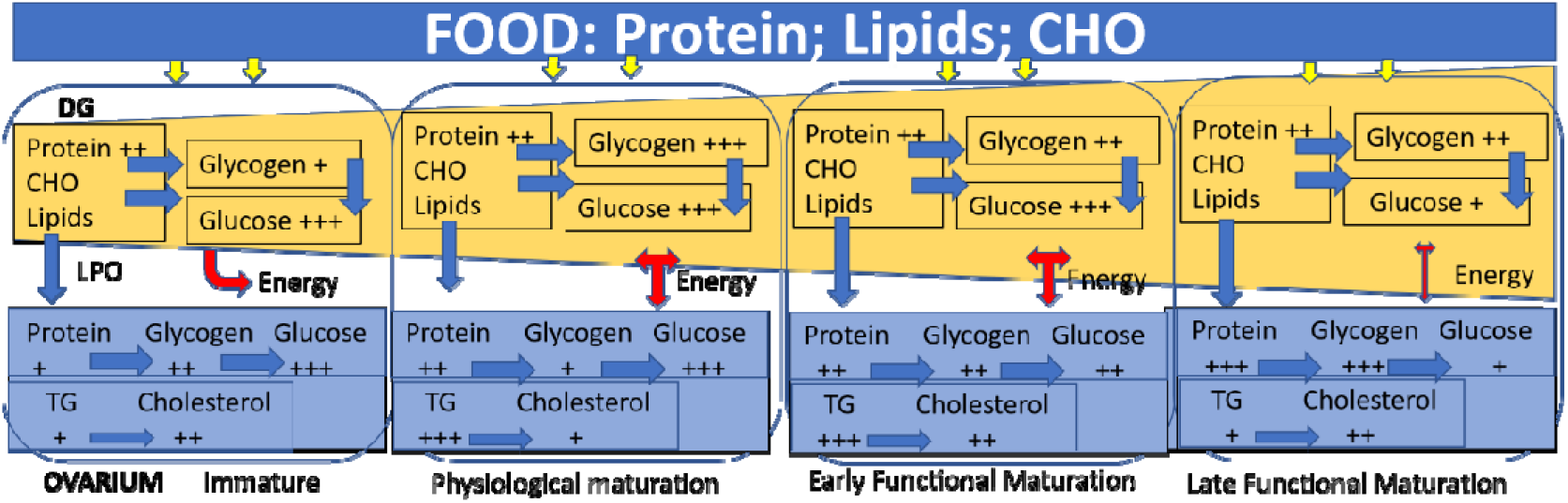
Physiological processes that occurs in digestive gland and ovarium during *O. mimus* female maturation. Nutrients, coming from the ingested food are processed into the digestive gland (DG) where proteins are used along maturation both as a source of carbohydrates via gluconeogenic pathway and to be transported as peptides and amino acids to internal tissues. Ingested carbohydrates (CHO) and glucose liberated from glycogen are used mainly as a source of energy into the DG and other tissues at the beginning of maturation and until the vitellogenesis start (early functional maturation) (Gallardo el al., 2017). In ovarium, protein and glycogen follow the same process than DG, supporting the glucose production until early functional maturation stage. Lipids (measured as triacyl glycerides: TG) and free cholesterol are absorbed in DG and transported by lipoproteins (LP) to ovarium, between other tissues (Heras & Pollero, 1990; Heras & Pollero, 1992). TG are accumulated during physiological maturation and early functional maturation and presumably converted at least in part, as cholesterol at the end of the maturation process (Estefanell *et al*., 2017).

The dynamics of glycogen and glucose observed in the ovarium of *O. mimus* suggest that there are biochemical regulatory mechanisms involved in storing and mobilization of nutrients. In the present study, high glucose levels were observed in the ovarium of immature females and during physiological maturation. This suggests that glucose could be used as a source of energy at the beginning of the complex processes involved in oocytes synthesis. However, a decrease of glucose levels was recorded during the functional maturation, with the lowest levels detected at the end of the process. If glucose is used only as a source of energy, why a reduction in glucose was recorded when the vitellogenesis process was at its maximum level? It is possible that the ovarium required less energy in the last part of the vitellogenesis than in previous maturation stages, or alternatively, glucose was reduced to avoid its inhibitory effects. Excess of glucose inhibits vitellogenin uptake in insects (Kunkel et al., 1987). It is therefore possible that a reduction of ovarium glucose levels in octopus could be associated with mechanisms to avoid physiological problems during the last part of the reproductive process. Glucose levels may thus be used only as a source of metabolic energy allowing the adequate vitellogenin uptake. If some mechanism of control exist, it also may be involved in glycogen synthesis and storing. During the late functional maturation, an increment of glycogen was registered in the ovarium. This suggests that those molecules were directly stored in the eggs used as a source of energy during embryo development and/or to maintain the physiological integrity of females during parental care that occurs after the spawning (Roumbedakis et al. 2017).

As was previously observed in *O. vulgaris* (Rosa et al., 2004), in *O. mimus* there was no apparent mobilization of lipids from the DG to the ovarium. In cephalopods, the nutrients channelled to the ovarium are thus likely obtained from the food and not from reserve tissues, as was observed in laboratory studies (Caamal-Monsreal et al. 2015; Rosa et al. 2005; Tercero-Iglesias et al. 2015).

In the present study, high levels of TG and cholesterol were detected in immature females, indicating that even before maturation the females used the ovary as a reserve of lipid. This means that nutrients come directly from the food and are not stored in muscle or the digestive gland (Rosa et al., 2005), and that the ovary itself is a reserve site for nutrients that will be used at least at the beginning of the maturation process (Fig. 9)

The increment of TG observed after the maturation processes started, and the subsequent reduction of cholesterol suggest that cholesterol was required during the physiological maturation (oocytes growth), probably as a structural component into the oocytes membranes. This process is characterized at the beginning by the formation of oocytes, which after growth will be transformed in secondary oocytes surrounded by the follicle cells without yolk (Avila-Poveda et al. 2016). It is highly likely that the oocytes and follicle cells synthesis require cholesterol during this process because it is an essential component of the biological membranes (Zubay 1983). High levels of TG were registered once the vitellogenesis started (early functional maturation process), indicating that fatty acids are also stored in the ovarium probably to be used in the yolk synthesis. This is supported by low levels of TG in the ovarium of the *O. mimus* females at the end of the maturation process (late functional maturation). Fatty acids accumulated as TG were likely transformed and stored into the eggs as yolk. Moderated cholesterol levels detected at the end of the maturation process suggest that cholesterol was also stored in the yolk to be used by the embryos throughout their development (Estefanell et al. 2017).

As expected, the biochemical and physiological processes in embryos are highly dynamics following two well identified developmental phases: organogenesis and growth (Boletzky 1987; Naef 1928). In many octopus species the first phase occurs between stage I to XII-XIII where the first embryo inversion allows the embryo to growth in the proximal side of the egg (Boletzky, 1987). During this phase, the nervous system is developed and the retina pigmentation is evident around stage X to complete organogenesis in *O. mimus* embryos at stages XII-XIII (Castro-Fuentes et al. 2002).

Our results indicate that soluble proteins and amino acids may be used as a source of glycogen, maintaining a stable and permanent supply of glucose along embryo development even during the growth phase. These results suggest that glucose is not the most important source of energy for embryos and that regulation of the gluconeogenic pathway works as a mechanism of the glucose supply. Regulation thus appears to be coupled with the phases of embryo development, without control until stage XI, and with control from stage XII onwards (Fig. 10). Our findings also suggest that energetic demands of embryos in the first phase of development were relatively low, without significantly mobilization of energetic substrates and its associated enzymes (Fig. 10). The molecules identified with redox stress and that were maintained in the eggs without changes suggest that the antioxidant defence mechanisms were inactive until the stage XIV.

**Fig. 10.**
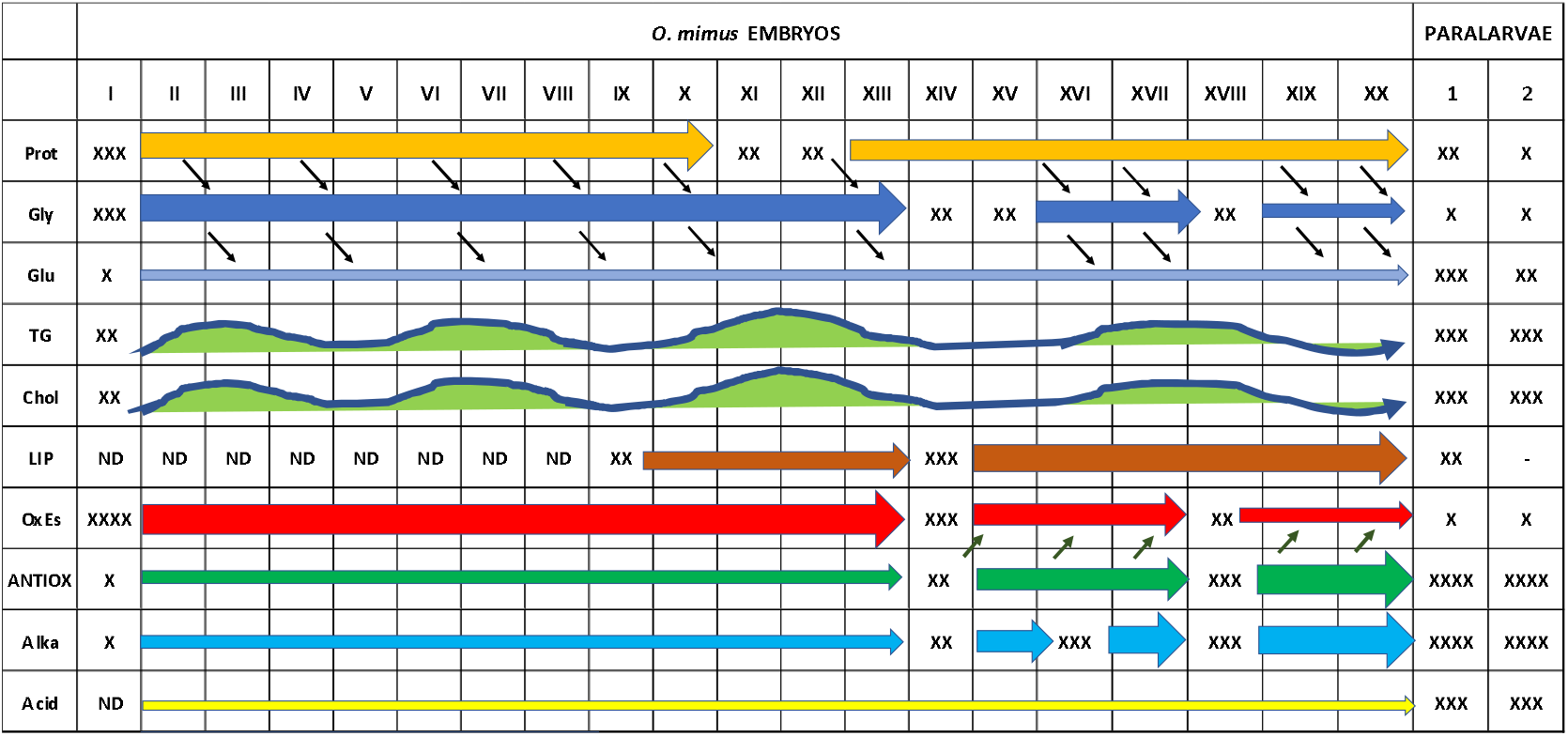
*O. mimus* biochemical and physiological dynamics through embryo development and after hatch. Gly: glycogen; Glu: glucose; Prot: soluble protein; TG: triacyl glycerides; Chol: Cholesterol; Alka: Alkaline enzymes; Acid: Acidic Enzymes; LIP: Lipases; OxEs: Oxidative stress; ANTIOX: Antioxidant defence mechanisms. Arrows indicate the possible metabolic pathway that are carried out into the embryos. Soluble proteins support the glycogen production via gluconeogenic pathways, while glycogen liberate glucose in a stable and permanent form through embryo development to maintain a stable and efficient metabolizable energy supply. Coming from the yolk, TG and Cholesterol are used in pulses while lipases activity was evident from the stage X onwards, when the organogenesis starts. At the beginning of the embryo development high levels of oxidative stress were observed, probably from residual ROS produced in the female ovarium. *O. mimus* embryos were able to start the elimination of ROS when they reached the stage XIV, when the activity of the antioxidant defence mechanisms started activities. Activities of alkaline and acidic digestive enzymes were evident at the end of embryo development and in hatchlings, when presumably the digestive gland is in preparation to ingest food.

The reduction on soluble protein and glycogen, and the increment of lipases activity in stage XIV embryos and onward indicates that the mobilization of yolk started at that stage (Fig. 10). Studies on embryos of *O. maya* and *O. mimus* found significant mobilization of yolk from stage XIV to stage XV suggesting that mobilization of nutrients from yolk marks the start of the embryonic growth (Caamal-Monsreal et al. 2016; Sánchez-García et al. 2017). It was observed that hearth beats in *O. mimus* increase with embryo development and reach a relative stability from stage XV, when the growth of embryos was evident (Warnke 1999). Power growth rate of *Enteroctopus megalocyathus* embryos was obtained from stage XII (Uriarte et al. 2016), suggesting that this phase of growth after organogenesis is a common characteristic among octopus species. The metabolic demand of cephalopods embryos increases with development as was observed in *Sepia officinalis, Loligo vulgaris* (Pimentel et al. 2012), *E. megalocyathus* (Uriarte et al. 2016), *O. vulgaris* (Parra et al. 2000), *O. mimus* (Uriarte et al. 2012), and *O. maya* (Caamal-Monsreal et al. 2016; Sánchez-García et al. 2017). Reactive oxygen species (ROS) are produced during this process, which consequently leads to oxidative stress (Regoli et al. 2011). To prevent oxidative stress and keep the balance of the cell aerobic organisms have evolved an efficient antioxidant defence system that consists of both non-enzymatic small antioxidant molecules (e.g. reduced glutathione (GSH), ascorbic acid (AA), carotenoids, etc.) and a cascade of enzymes (e.g. superoxide dis-mutase (SOD), catalase (CAT), and glutathione peroxidase (GPx) (Regoli & Giuliani 2014). A recent study on the role of the enzymatic antioxidant system in the thermal adaptation of *O. vulgaris* and *O. maya* embryos suggests that early developmental stages of cephalopods have temperature-regulated mechanisms to avoid oxidative stress (Repolho et al. 2014; Sánchez-García et al. 2017). In the present study, it was observed that ROS of *O. mimus* embryos were almost totally eliminated during the growth phase of the embryo development due to the activation of the antioxidant defence mechanisms in stage XIV, indicating the coupling between metabolic demands and the functioning of the antioxidant defence system against oxidant stress.

During the burst of anaerobic swimming of *O. vulgaris* paralarvae the energy is obtained from glucose and from the Arginine phosphate system mediated by lactate dehydrogenase (LDH) and octopine dehydrogenase (ODH) respectively (Morales et al. 2017). Both systems require pyruvate, either by the gluconeogenic route or via the degradation of amino acids by transamination (Zubay 1983). We hypothesize that the operating mechanism in *O. mimus* embryosmay be associated with amino acids and lipid catalysis because both substrates (mainly arginine and glycerol) are involved in pyruvate production in cephalopods (Morales et al. 2017). The apparent lack of activity of lipases until stage X, and the moderate activity of alkaline and acidic enzymes along the development of *O. mimus* embryos may be involved in physiological regulation of the energy supply in those organisms. Accordingly, the yolk consumption in *O. maya* embryos starts after the stage XIV, when organogenesis ends (Caamal-Monsreal et al. 2015; Sánchez-García et al. 2017) and the antioxidant defense mechanisms start its activity.

## Conclusions

This study demonstrates that ovarium can be used as a reserve of some nutrients for reproduction. Acyl glycerides where stored at the beginning of the maturation processes followed by cholesterol. Acyl glycerides and cholesterol were energetically supported by glucose and derived from glycogen following gluconeogenic pathways. This suggests that a control mechanism of protein-glycogen-glucose may be operating in *O. mimus* ovarian. We hypothesize that glucose is the energetic support at the beginning of ovarian maturation and also has the potential role as inhibitor of vitellogenin uptake at the end of the processes. It was observed that embryos during organogenesis, nutrients and enzymes (metabolic, digestive and REDOX system) were maintained without significant changes and in a low activity. During organogenesis, yolk was maintained constant indicating that blastulation and gastrulation do not appear to be influenced by the size of the yolk mass (Boletzky 2003). In this phase, the outer yolk sac envelope combined with its role of reserve of nutrients have the provisional function of gill and heart. Its activity is supported by a dense ciliature covering the entire yolk sac surface that maintain perivitelline fluids in circulation and oxygen uptake by the embryo (Boletzky 1989). Our results suggest that this activity has a low energetic cost. In contrast, it was observed that during the embryo growth, when the activity of the hearth was evident (Castro-Fuentes et al. 2002), there waas mobilization of nutrients and activation of the metabolic and digestive enzymes, as well as increments in the consumption of yolk and glycogen, and the reduction in molecules associated with oxidative stress. This allowed paralarvae to hatch with the antioxidant defence mechanisms ready to support the ROS production.

